# Moderately acid-tolerant *Nitrospira* strains obtained by single-colony isolation on plates

**DOI:** 10.64898/2026.11.10.765609

**Authors:** Yueyue Zhu, Yangyang Peng, Yuyi Zheng, Weitie Lin, Jianfei Luo

## Abstract

*Nitrospira* are difficult to cultivate, and pure cultures are mainly obtained by dilution-to-extinction combined with other methods, a process that usually takes years to over a decade. Here, we isolated three *Nitrospira* strains from activated sludge and sediment within one year using conventional single-colony isolation. Pure cultures of two strains could grow and reform single colonies within 25-60 days on all three types of plates prepared with agar, agarose, and gellan gum. Phylogenetic and phylogenomic analyses placed them within lineage I and indicated that they represent potential novel species, with the provisional names ‘*Nitrospira monocolonialis*’ for strain Z02 and ‘*Nitrospira limosa*’ for strains Z47 and Z70. All three strains were moderately acid-tolerant, with a minimum tolerable pH of 5.0; strain Z02 showed the highest acid tolerance, retaining 74% of its activity at pH 5.5. Strain Z02 was also highly adaptable to oxygen concentrations of 1-21%, retaining 78% and 85% of its maximum growth rate at 1% and 21% O_2_, respectively. Compared with the representative lineage I strain *Nitrospira defluvii*, all three strains contain superoxide dismutase encoding genes and some unique genes. Comparative transcriptomic analysis suggested that strain Z02 upregulated genes involved in stress responses, cell envelope modification, and putative type IVa pilus systems under acidic condition. Overall, this study provides the first demonstration that *Nitrospira* can grow on plates prepared with common gelling agents and form single colonies within a short period, and that *Nitrospira* can also tolerate moderately acid conditions.

## Introduction

Members of the genus *Nitrospira* are widely distributed in natural and engineered ecosystems and contribute substantially to the biogeochemical cycling of nitrogen. The genus includes canonical nitrite-oxidizing bacteria (NOB), which oxidize nitrite to nitrate, and complete ammonia oxidizers (comammox), which oxidize ammonia to nitrate within a single organism [1–4]. Since *Nitrospira marina* [5] was first isolated, cultured representatives have expanded to include *N. moscoviensis* [6], *N. japonica* [7], *N. defluvii* [8], *N. lenta* [8], *N. inopinata* [3, 9] and *N. tepida* [10], as well as numerous enrichment cultures (https://lpsn.dsmz.de). As difficult-to-cultivate bacteria, isolation and purification of *Nitrospira* strains often takes several years or even more than a decade [1]. For example, *Candidatus* (*Ca.*) Nitrospria bockiana took 12 years [11], *N. defluvii* took 4 years to reach pure culture from an 86% enrichment [8], and a *N. marina*-like took 4 years [12]. These cultures were obtained mainly through dilution-to-extinction, density-gradient centrifugation, selective antimicrobial treatment, micromanipulation, and microcolony sorting, or combinations thereof (Fig. S1). Dilution-to-extinction has been used for nitrifier isolation for over 130 years and remains widely used [13].

Compared with these methods, single-colony isolation on conventional solid plates, the most used and effective approach for microbial isolation, has rarely been reported for *Nitrospira*. During isolation of *N. defluvii*, a highly enriched culture formed colonies with heterotrophic bacteria on mixotrophic agarose medium; however, no pure culture was obtained by single-colony isolation, and the purified culture did not reform colonies under the conditions tested [8]. More recently, common gelling agents, including agar, agarose and gellan gum, were found to contain toxic substances inhibiting diverse nitrifiers in liquid culture and on solid plates; removal of these inhibitors using washed gellan gum, but not agar or agarose, enabled the tested nitrifiers including *Nitrospira* sp. ND1, *N. japonica* J1 and *Nitrospira* sp. KM1 to form single colonies [14]. In addition, *Nitrospira* traits such as extremely low growth rate, cell aggregation, and close interactions with heterotrophs make single-colony isolation difficult [7, 15, 16].

During two-step nitrification, ammonia oxidation releases protons and decreases pH; therefore, NOB require acid tolerance to cooperate with ammonia oxidizers such as ammonia-oxidizing bacteria (AOB) and ammonia-oxidizing archaea (AOA) and coexist effectively. Several acid-tolerant or acidophilic NOB have been found to grow or oxidize nitrite at pH 3.5–5.5, including pure cultures of *Nitrobacter* sp. Io acid [17], *Nitrobacter* sp. A67 [18], “*Ca.* Nitrobacter laanbroekii” NHB1 [19] and *Nitrobacter* sp. JJSN [20], as well as enrichment cultures of “*Ca.* Nitrocaldera robusta” [21] and “*Ca.* Nitrotoga” sp. CP45 [22]. In contrast, most characterized strains of canonical *Nitrospira* grow optimally or are maintained at neutral to mildly alkaline pH, with weak growth or nitrite oxidation below pH 6.0 [5, 6, 23, 24]. Cultivation evidence for acid-tolerant *Nitrospira* comes only from enrichment cultures [25–28], with limited pure-culture characterization. To date, *Nitrospira*-related populations are widely detected in acidic natural and engineered environments, including nitrifying biofilms [29, 30], sediments [31], agricultural soils [32] and mine lakes [33], and usually show higher abundances than *Nitrobacter*-related groups [34–36]. These observations suggest that many *Nitrospira* strains exhibit acid tolerance similar to that of *Nitrobacter* and drive nitrification with ammonia oxidizers in acidic environments.

The study aimed to: (i) isolate *Nitrospira* strains from different environments by single-colony isolation using conventional solid plate and demonstrate that plates made with common gelling agents can potentially be used for single-colony isolation and cultivation of *Nitrospira*; (ii) demonstrate using pure cultures that *Nitrospira* can tolerate acidic conditions; and (iii) explore possible acid-tolerance mechanisms of *Nitrospira* at the omics level. This study ultimately obtained three *Nitrospira* strains by single-colony isolation and demonstrated at the cellular level that they are moderately acid-tolerant.

## Materials and Methods

### Sampling and enrichment cultivation of Nitrospira

Activated sludges were collected from wastewater treatment plants in Foshan (FS) and Guangzhou (GZ); sediment was collected from a ditch in Shantou (ST), all in Guangdong Province, China. The FS was first enriched for comammox with ammonium as the sole energy source. For *Nitrospira* NOB enrichment, a 1% (v/v) comammox culture from FS, 1 g sludge from GZ, or 1 g sediment from ST was inoculated into a 125-mL serum bottle (butyl rubber stopper, metal crimp top) containing 50 mL of basal medium. The basal medium consisted autoclavable components: NaCl (1.0 g/L), CaCl₂·2H₂O (0.02 g/L), and MgSO₄·7H₂O (0.05 g/L); and filter-sterilized (0.22-µm) components: K₂HPO₄ (0.653 g/L), KH₂PO₄ (0.17 g/L), trace element stock solution (1 mL/L), and NaNO₂ and NaHCO₃ at condition-specific concentrations. Trace element stock solution (per L) contained FeCl₂·4H₂O 1.5 g, CoCl₂·6H₂O 190 mg, MnCl₂·4H₂O 100 mg, ZnCl₂ 70 mg, H₃BO₃ 62 mg, Na₂MoO₄·2H₂O 36 mg, NiCl₂·6H₂O 24 mg, and CuCl₂·2H₂O 17 mg. Two-stage fed-batch cultivation was performed with nitrite at 10 μM (stage 1) and 50 μM (stage 2) (Fig. 1A).

**Figure 1.**
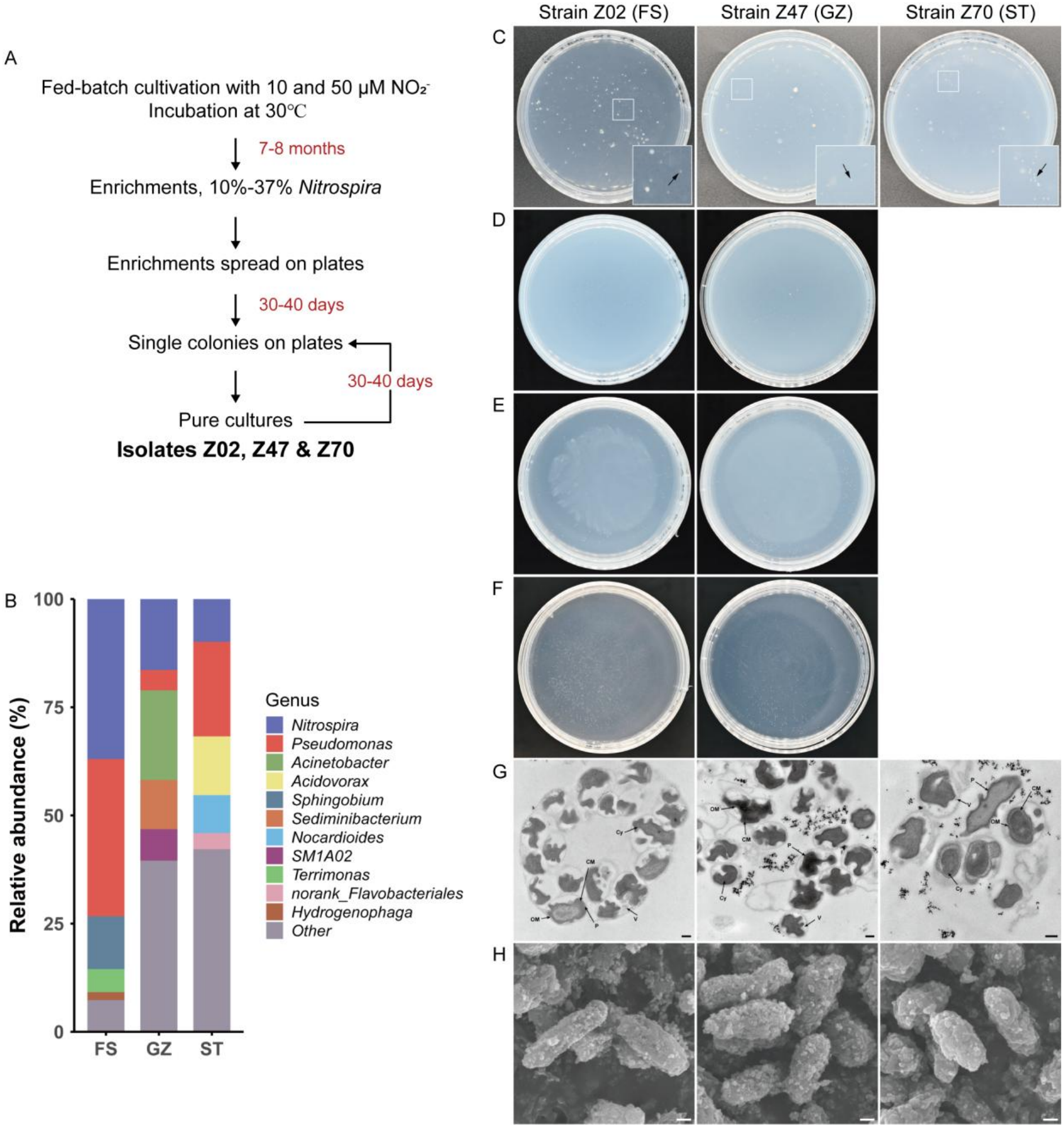
Enrichment, single colony isolation and morphology of *Nitrospira* strains. **A)** Enrichment workflow. **B)** Genus-level community composition based on full-length 16S rRNA amplicon sequencing. The five most abundant genera in samples are shown; all others are grouped as “Other”. **C)** Colonies of enrichment cultures grown on agarose plates after inoculation for 2 months. *Nitrospira* colonies are indicated by black arrows. **D-F)** Colonies of pure cultures grown on agarose plates **(D)**, agar plates **(E)** or gellan gum plates **(F)** after inoculation for 2-3 months. The granular or banded precipitate background in agar and gellan gum plates resulted from the preparation process. **G)** Transmission electron micrographs of *Nitrospira* cells. Black arrows indicate the cytoplasm (Cy), cytoplasmic membrane (CM), periplasm (P), outer membrane (OM), and membrane vesicles (V). The scale bar represents 0.2 μm. **H)** Scanning electron micrographs of *Nitrospira* cells. The scale bar represents 0.2 μm.

Feeding was performed 5-7 times per stage, with NaHCO₃ added at one-tenth the molar amount of nitrite. After each round, the culture was transferred at a 0.4% (v/v) inoculum into the next round, and this serial transfer was repeated 6-7 times. Cultivations were at 30 ℃ in the dark without agitation, with pH maintained at 7.2–7.4 throughout enrichment.

### Single-colony isolation and cultivation of Nitrospira strains on plates

When *Nitrospira* abundance exceeded 10%, enrichment cultures were subjected to single-colony isolation using 1.2% (w/v) agarose as gelling agent. Briefly, enrichment cultures were harvested when approximately 80% of supplied nitrite was consumed, serially diluted tenfold, and 100 μL of each dilution was spread onto agarose plates; each plate was sealed with polyethylene film, enclosed in polyethylene bags, and incubated at 30 ℃ in the dark. To prepare solid plates, agarose was autoclaved together with the autoclavable basal medium components. After cooling to 50 ℃, the following filter-sterilized components were added: K₂HPO₄ (0.653 g/L), KH₂PO₄ (0.17 g/L), NH₄Cl (3 mM), NaHCO₃ (3 mM), NaNO₂ (10 mM), and trace element solution (2‰, v/v). After 30-40 days, individual colonies were picked and transferred to liquid basal medium to confirm nitrite-oxidizing activity. Positive cultures were inoculated into Luria-Bertani or R2A liquid medium and simultaneously spread onto corresponding agar plates to check purity. Purity was also verified by 16S rRNA gene sequencing; if not pure, the culture was re-spread onto agarose plates and individual colonies were re-streaked until a pure culture was obtained.

To demonstrate that pure cultures of *Nitrospira* could reform colonies on solid plate prepared with common gelling agents, agar and agarose plates were prepared with 1.2% (w/v) agar or agarose as above. Gellan gum plates used a modified two-component procedure: solution A (NaCl 1.0 g/L, CaCl₂·2H₂O 0.1 g/L, MgSO₄·7H₂O 0.25 g/L) and solution B (same salts in solution A plus 1.6% (w/v) phytagel; Sigma-Aldrich, P8169) were autoclaved separately; after cooling to 50 ℃, filter-sterilized components were added to solution A at 2× final concentrations, then equal volume of solution B was added and mixed thoroughly.

Diluted culture was spread onto plates, sealed as above, and incubated at 30 ℃ in the dark.

### Full-length 16S rRNA gene amplicon sequencing

DNA from enrichment cultures was extracted using a modified CTAB method [37] and subjected to 16S rRNA gene amplicon sequencing on the PacBio Sequel platform by Sangon Biotech (Shanghai, China) with primer pair 27F (5′-AGRGTTYGATYMTGGCTCAG-3′) and 1492R (5′-RGYTACCTTGTTACGACTT-3′). Raw reads were processed into circular consensus sequences (CCS) through self-correction and demultiplexed by barcode. Amplicon sequence variants (ASVs) were inferred using the DADA2 pipeline v1.14.0 [38] and taxonomically assigned against the nt (202410) database using Blast+ v2.10.0 [39].

### Whole-genome sequencing, comparative genomic and phylogenetic analyses

*Nitrospira* cells were harvested by centrifugation at 10,000 × *g* for 10 min, and genomic DNA was extracted using the E.Z.N.A.® Bacterial DNA Kit (OMEGA, D3350, USA). Whole-genome sequencing was performed by Shanghai Majorbio Bio-pharm Technology Co., Ltd. (Shanghai, China) using a hybrid strategy combining the PacBio Sequel IIe and the Illumina NovaSeq 6000 platforms. Data were analyzed on Majorbio Cloud Platform [40]. Genome completeness and contamination were estimated with CheckM2 [41] using default parameters. Protein-coding genes were predicted with Prodigal v2.6.3 [42] and annotated with DIAMOND v0.9.24 [43] against Swiss-Prot and TrEMBL databases (release 202602) [44]. Genomes were also annotated with Bakta v1.11.0 [45] (full database v6.0) and the NCBI PGAP [46].

Comparative genomic analysis used strains Z02, Z47 and Z70, plus selected genomes and metagenome-assembled genomes representing distinct *Nitrospira* lineages (Table S1). OrthoANIu [47] and EzAAI v1.2.4 [48] were used to calculate average nucleotide identity (ANI) and average amino acid identity (AAI), respectively. Maximum-likelihood (ML) and Bayesian trees based on 16S rRNA gene and *nxrB* sequences were inferred with IQ-TREE v3.1.2 [49–51] and MrBayes v3.2.7 [52]. IQ-TREE used-m MFP - B 1000. MrBayes used GTR+Γ4 for the 16S rRNA gene. The *nxrB* analyses used either an unpartitioned GTR+I+Γ4 model or codon partitions with independent GTR+Γ4 parameters and variable partition rates. Each analysis ran 5,000,000 generations with four chains, discarding the first 25% as burn-in. For phylogenomics, single-copy orthologues were identified with OrthoFinder v3.1.4 [53] using-M dendroblast, aligned, trimmed, concatenated, and used for ML inference in IQ-TREE using the above settings. Trees and ANI/AAI heatmaps were visualized in R v4.5.1; other trees were annotated in iTOL v7.4.1 [54].

### Morphological and physicochemical characterization of Nitrospira isolates

Cells in exponential growth phase were harvested by centrifugation at 6,000 × *g* for 10 min at 10 ℃, washed three times with 10 mM phosphate buffer (pH 7.0), and fixed with 2.5% (w/v) glutaraldehyde at 4 ℃ for 12 h. For transmission electron microscopy (TEM), fixed cells were rinsed with 0.1 M phosphate buffer (pH 7.4), pre-embedded in 1.5% (w/v) agarose, post-fixed with 1% (w/v) osmium tetroxide in the same buffer at room temperature for 2 h, dehydrated in a graded ethanol series (30–100%, v/v), embedded in Embed 812 resin, stained with 1% (w/v) lead citrate, and observed under a Hitachi HT7800 electron microscope. For scanning electron microscopy (SEM), fixed cells were dehydrated in a graded ethanol series (30–100%, v/v), replaced with tert-butanol, vacuum freeze-dried, sputter-coated with gold using an EMS 150TES sputter coater, and observed under a Zeiss Merlin electron microscope.

Physicochemical analyses were performed in 125-mL serum bottles containing 50 mL of medium and sealed with butyl rubber stoppers. Conditions tested: (i) 1–100 mM initial nitrite with a 1% (v/v) inoculum and 2.5 mM NaHCO₃; (ii) 20–40 ℃; (iii) 1–21% (v/v) oxygen in the headspace; (iv) pH 5.0–9.0; and (v) combined pH 4.5–5.5 and 0.03–1 mM initial nitrite. Unless otherwise specified, cultures were inoculated at 0.1% (v/v) inoculum and supplemented with 0.5 mM NaHCO₃ and 1 mM nitrite. Incubations were performed at 30 ℃ in the dark without agitation. Initial pH was stabilized with 5 mM MES, HEPES, or Tris adjusted with NaOH or HCl.

Oxygen concentrations in the headspace were established and measured as described previously [55]. Nitrite and nitrate concentrations were measured using a vanadium chloride reduction-based Griess assay adapted from previously described protocols [56, 57]. For nitrite determination, 100 μL of sample was mixed with 50 μL of diazotizing reagent (0.2%, w/v, sulfanilamide in 10%, v/v, HCl) and 50 μL of coupling reagent [0.1%, w/v, N-(1-naphthyl) ethylenediamine in water], and the absorbance was measured at 540 nm. For nitrate determination, 100 μL of VCl₃ solution (0.8%, w/v, in 1 M HCl) was added to the same mixture, incubated at 50 ℃ in the dark for 30 min, and measured at 540 nm. Nitrate was obtained by subtracting initial nitrite from measured total. As described previously, nitrate concentrations during exponential growth were fitted to an exponential model to estimate the apparent maximum specific growth rate (μ_max_, _app_) (Fig. S5A) [19].

### Kinetic analysis of nitrite oxidation

The nitrite oxidation kinetic assay was adapted from a previous method [58]. Briefly, *Nitrospira* cells grown to the pre-stationary-phase were collected by centrifugation at 6,000 × *g* for 10 min at 10 ℃, washed twice with equal volumes of culture medium lacking NaHCO₃ and nitrite, resuspended to the original volume, and dispensed into serum tubes sealed with butyl rubber stoppers. Incubations were at 30 ℃ in the dark with shaking at 200 rpm in a water bath shaker. Tubes were pre-incubated 30 min to deplete endogenous nitrite. Sodium nitrite was then added to the serum tubes at concentrations of 10–1,500 μM to initiate nitrite oxidation. After incubation for 10–60 min, samples were taken, immediately filtered through 0.22-µm filters, and filtrates were incubated at 90 ℃ for 10 min in a water bath to inactivate the cells. The oxidation rate at each initial nitrite concentration was calculated from the difference in nitrite before and after incubation, and apparent kinetic parameters were obtained by fitting to the Michaelis–Menten equation. To normalize the maximum specific nitrite oxidation rates (V_max_) to unit cell biomass, a 40 mL seed culture prepared for kinetic analysis was harvested by centrifugation at 12,000 × *g* for 10 min at 4 ℃, cells were lysed by hot alkaline lysis method [59], and protein content was determined using the Pierce BCA Protein Assay Kit (Thermo Scientific, A65453, USA) following the manufacturer’s instructions.

### Comparative transcriptomic analysis

Strain Z02 was cultivated in basal medium and MES-buffered media described above, with pH maintained at 7.4 and 5.5, respectively. Cells grown to exponential phase were harvested by centrifugation at 12,000 × *g* for 5 min at 4 ℃. Total RNA was extracted using RNAiso Plus Reagent (Takara, 9108, Japan) according to the manufacturer’s instructions. Transcriptome sequencing was performed by Shanghai Majorbio Bio-pharm Technology Co., Ltd. (Shanghai, China) on the Illumina NovaSeq X Plus platform. Reads were analyzed on Majorbio Cloud Platform [40] using the complete genome of strain Z02 as reference. Putative pilus and secretion systems were identified using MacSyFinder v2.1.6 [60] with the TXSScan models [61].

### Statistical analysis

Normality and homogeneity of variance were evaluated using the Shapiro–Wilk and Levene tests, respectively. Group means were compared using Welch’s ANOVA followed by the Games–Howell post hoc test. P < 0.05 was considered statistically significant.

## Results and discussion

### Enrichment cultivation and single-colony isolation of Nitrospira strains

After 7-8 months of cultivation, full-length 16S rRNA gene amplicon sequencing showed that *Nitrospira* reached a relative abundance of 10-37%, with no other nitrifiers detected in the three enrichment cultures (Fig. 1B). PCR analysis also showed that AOB, AOA and *Nitrobacter* were undetectable (data not shown). The FS sample had the highest abundance and the simplest bacterial community, likely due to its origin from a well-enriched comammox culture. However, the comammox that accounting for > 10% of the initial seed culture was absent, probably because comammox *Nitrospira* cannot assimilate nitrite as a nitrogen source [62–64] and the medium lacked ammonium. ASV analysis showed that ASV2 was the only *Nitrospira*-related ASV in the FS sample, accounting for 37.0% of the community (Fig. S2A). The GZ sample also contained one *Nitrospira*-related ASV (ASV47), accounting for 16.4% of the community, whereas the ST sample contained ASV70 and ASV87, which accounted for 9.2% and 0.7% of the community, respectively (Fig. S2A).

The enrichment cultures were spread onto agarose plates. After 30-40 days, colonies typical of heterotrophic bacteria and translucent, pinpoint-like colonies appeared. Although colony distribution was irregular, both large and small colonies formed clear single colonies at 10⁻³ dilutions (Fig. 1C). Since *Nitrospira* single colony formation on agar or agarose plates had not been reported, the identity of these pinpoint colonies was initially uncertain. Their morphology and color matched those of the initial AOB colonies on agar or agarose plates in our pervious observations [55, 65] and of colonies formed by strains *Nitrospira* sp. ND1 and *N. japonica* on washed gellan gum plates [14]. These translucent, pinpoint colonies were picked and inoculated into liquid medium. All isolates showed nitrite-oxidizing activity with near-stoichiometric nitrite oxidation and nitrate production. Purity checks showed that more than half of the picked colonies were free of heterotrophic bacteria. 16S rRNA and *nxrB* genes sequencing showed single peaks in the nucleotide sequence chromatograms, and the sequences were closely related to *Nitrospira*, confirming pure cultures. Sequence alignment further showed that the 16S rRNA gene of these pure cultures showed 100% similarity with ASV2, ASV47, and ASV70, respectively. The isolates from FS, GZ, and ST samples were therefore designated *Nitrospira* strains Z02, Z47 and Z70, respectively.

### Re-formation of single colonies by Nitrospira strains on plates

Pure cultures of strains Z02, Z47 and Z70 were spread onto plates prepared with agarose, agar, or gellan gum. Strains Z02 and Z47 formed visible colonies on agarose plates only after 25 and 28 days, respectively, whereas strain Z70 did not form colonies even after > 3 months (Fig. 1D). After steaking a single colony, strain Z02 also formed visible single colonies on agarose plates after 35 days (Fig. S2B). With prolonged incubation time, colonies continued to enlarge; after 13 months, strain Z02 colonies reached approximately 2 mm in diameter (Fig. S2C-K). Strains Z02 and Z47, but not strain Z70, could also reform single colonies on agar and gellan gum plates (Fig. 1E, F). Apart from the time required, colony color and morphology on agar and gellan gum plates did not differ from those on agarose plates. Visible colonies of strains Z02 and Z47 formed after 28 days on agar plates, whereas on gellan gum plates they became visible after 37 and 61 days, respectively.

These results indicate that conventional solid plates are also suitable for isolating and purifying *Nitrospira* strains, but colony-forming ability differs among strains. Certain nutritional requirements or environmental adaptions may affect growth on plates. For example, toxic substances in gelling agents or produced during autoclaving could inhibit growth and colony formation [14]. Strain *N. defluvii* formed colonies on agarose plates only with heterotrophic bacteria [8], suggesting dependence on these heterotrophs for specific nutrients or physiological support. Similar patterns occurred in AOB cultivation: some AOB strains are highly oxygen-sensitive and cannot grow at relatively high oxygen concentrations either in liquid culture or on agar plates [55], and Mg^2+^ inhibition has also been reported, with removal of Mg^2+^ supporting AOB growth [65]. Therefore, identifying the underlying substances or mechanisms and removing toxic substances from gelling agents could enable isolation of more *Nitrospira* strains by single-colony isolation.

### Morphological observation of Nitrospira strains

*Nitrospira* strains Z02, Z47, and Z70 had similar cellular morphologies. Although aggregated, the cells had an enlarged periplasmic space and were separated by thick layers of surrounding extracellular polymeric substances (EPS) (Fig. 1G, H), a feature also observed in *N. moscoviensis* [6], *N. japonica* [7], and *N. defluvii* [8]. Owing to thick EPS covering, the cells did not show the typical spiral morphology; SEM showed that they appeared more like rod-shaped cells, 0.2–0.3 μm wide and 0.8–1.2 μm long (Fig. 1H). As *Nitrospira* are typically embedded in EPS, cell morphology is often difficult to observe visually, and SEM observations are rare; for example, no cell morphology could be discerned in SEM of *N. japonica* [7], whereas comammox *N. inopinata* showed a coiled spiral morphology [9].

Extensive EPS synthesis and cell embedding may be common in *Nitrospira* and could protect key enzymes of the reductive TCA (rTCA) pathway from oxygen toxicity. Unlike *Nitrospira*, *Nitrobacter* strains exhibit clear, clean cell morphologies and preferentially synthesize intracellular polyhydroxybutyrate (PHB) granules to store fixed carbon rather than EPS [66–68], possibly associated with their use of the oxygen-insensitive Calvin–Benson–Bassham (CBB) pathway for carbon fixation. EPS synthesis versus intracellular PHB accumulation may differ among NOB genera, but extracellular release of dissolved organic carbon (DOC) is common to NOB and other nitrifiers [69, 70]. Released organic matters can recruit heterotrophic bacteria, thereby alleviating the toxicity from factors such as Mg^2+^ or oxygen [65, 71]. Whether EPS is synthesized or DOC is released, these characteristics make heterotrophic bacteria difficult to remove during enrichment cultivation and are key factors limiting the isolation and purification of *Nitrospira* strains.

### Phylogenetic and phylogenomic analyses of Nitrospira strains

The genomes comprised complete circular chromosomes of 4.15, 4.43 and 4.32 Mb for strains Z02, Z47 and Z70, respectively, with G+C contents of 59.0-60.2%. Strains Z47 and Z70 additionally carried plasmids of approximately 0.19 and 0.22 Mb, respectively. CheckM2 estimated genome completeness at 99.97–100%, with contamination estimates of 0.58-3.49% (Table S1).

Maximum-likelihood (ML) analysis of 16S rRNA gene sequences placed all three strains within *Nitrospira* lineage I, in the same clade as *N. defluvii* (Fig. 2A), consistent with the Bayesian inference (BI) tree (Fig. S3). ML and unpartitioned BI trees from *nxrB* showed broadly similar topologies, whereas the codon-partitioned BI tree differed at several deep nodes (Fig. S4A–C). None of the three *nxrB* analyses reproduced the previously reported broad correspondence between lineage-level structures of *nxrB* and 16S rRNA gene trees within *Nitrospira* [72]. In all three *nxrB* trees, only strains Z47 and Z70 remained in a local clade with *N. defluvii*, whereas the two *nxrB* copies of strain Z02 formed a separate, deep branch outside this group. This branching reflected differences in NxrB similarity among the three strains and other *Nitrospira* strains: the NxrB of strain Z02 showed 98.6% similarity to *N. defluvii* and 98.1% to strains Z47 and Z70, whereas strain Z47 and Z70 both showed 99.5% similarity to *N. defluvii*. Apart from the most similar sequences, NxrB similarities to most other strains, including those from different lineages, differed by less than 2%. Consequently, NxrB-based phylogeny, unlike 16S rRNA gene phylogeny, does not show strong conservation among branches.

**Figure 2.**
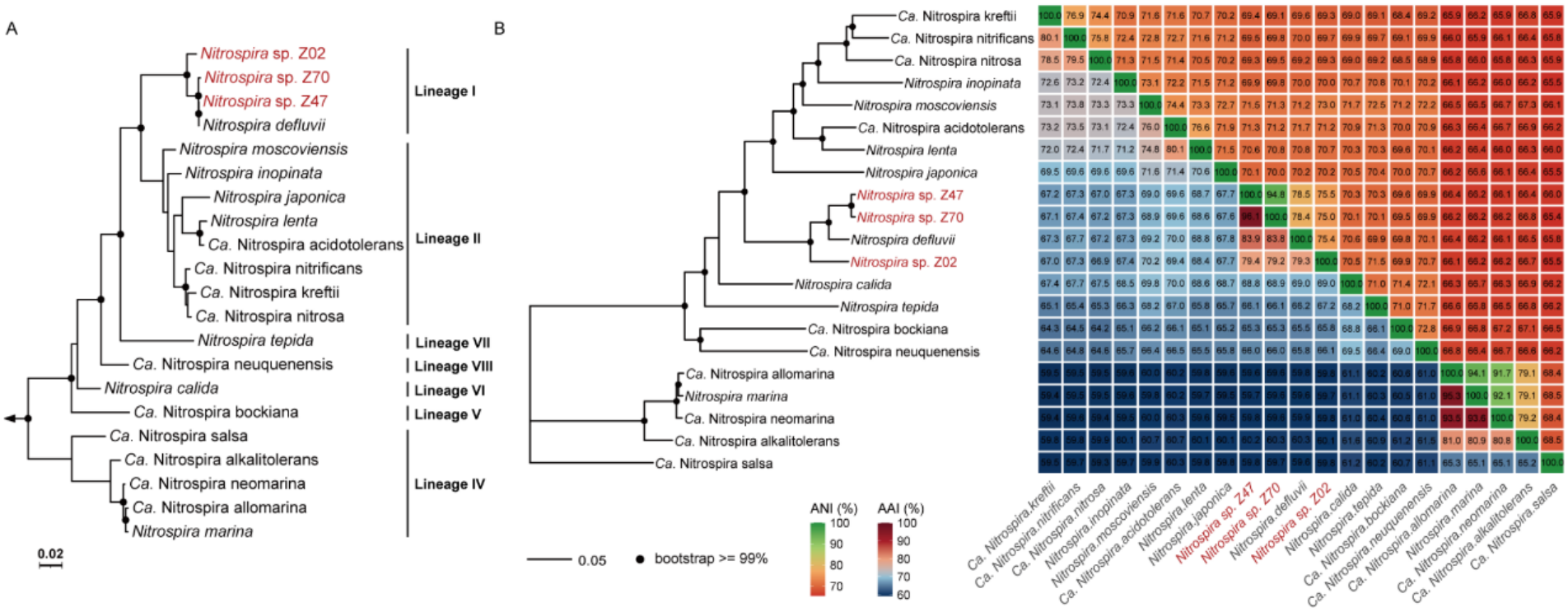
Phylogenetic and phylogenomic analyses of *Nitrospira* strains. **A)** Maximum-likelihood phylogenetic tree based on 16S rRNA gene sequences. Full-length sequences were used, except for the 976-bp partial sequence of “*Ca. Nitrospira* neuquenensis”. A total of 1,537 nucleotide sites were retained after alignment and trimming. Three *Leptospirillum* sequences from the family Nitrospiraceae were used as the outgroup, indicated by an arrow. Black dots denote branches with ultrafast bootstrap support ≥95%. Sublineages of the genus *Nitrospira* were indicated. **B)** Maximum-likelihood phylogenetic tree based on whole-genome sequences and heatmap of ANI and AAI values. The phylogenetic tree was inferred from 934 single-copy orthologues, with 290,403 amino acid sites retained after alignment, trimming, and concatenation. The upper and lower triangles of the heatmap correspond to ANI and AAI values, respectively.

Strain Z02 shared 98.25% and 98.18% 16S rRNA gene similarities with strains Z47 and Z70, respectively. Among recognized species, strain Z02 showed the highest similarity to *N. defluvii* (98.25%), below the 98.65% species threshold [73]; strains Z47 and Z70 were also most similar to *N. defluvii*, with similarities of 100% and 99.81%, respectively. A genome-scale ML tree based on concatenated single-copy orthologues supported the phylogenomic placement of all three strains (Fig. 2B). Strains Z47 and Z70 shared an ANI of 94.8%, whereas their ANI to strain Z02 were 75.5% and 75.0%, respectively. All ANI values between the three strains and described *Nitrospira* species were below 80%, well below the 94-96% species boundary [74, 75]. Strains Z47 and Z70 shared an AAI of 96.1% and showed 79.4% and 79.2% AAI to strain Z02, respectively. Their AAIs to *N. defluvii*, were 79.3%, 83.9% and 83.8%, respectively, whereas AAIs to other strains were below 70%, well below the 95% species threshold [76].

Based on phylogenetic and phylogenomic analyses, together with subsequent phenotypic analyses, they represent potential novel species within genus *Nitrospira*. We provisionally name strain Z02 as ‘*Nitrospira monocolonialis*’ (mo.no.co.lo.ni.a’lis. Gr. pref. *mono-*, single; L. fem. n. *colonia*, colony; N.L. masc. adj. *monocolonialis*, pertaining to a single colony / forming single colonies), and strains Z47 and Z70 as ‘*Nitrospira limosa*’ (li.mo’sa. L. fem. adj. *limosa*, slimy, referring to activated or sediment sludge).

### Genome-based metabolic potential of Nitrospira strains

Genome annotation showed that strains Z02, Z47 and Z70 shared a conserved genetic basis for nitrite oxidation and autotrophic carbon fixation (Fig. 3; Tables S2–S5). Each genome contains two sets of *nxrAB* genes and three independently located *nxrC* genes. All three genomes harbor genes encoding assimilatory nitrite reductase NirA, copper-containing nitrite reductase NirK, cyanase CynS, multiple cytochrome bd-like terminal oxidases, cytochrome c maturation system II, and a group 4 hydrogenase-related complex comprising HyfBCEFGI homologues (Fig. 3; Tables S2–S5). Each genome also contains a complete formate-associated module with genes encoding formate transporter FocA and soluble formate dehydrogenase genes FdsABC, suggesting potential formate oxidation capacity. Formate oxidation genes occur in *Nitrospira* lineage I strains and *N. moscoviensis* (lineage II), but activity has been demonstrated only in strain *N. moscoviensis* [77]. Except for *N. lenta*, other *Nitrospira* NOB genomes harbor genes encoding group 2a, 3b, or 4f-like hydrogenases, or two or three of these types (Table S2), although hydrogen-oxidizing activity is known only in *N. moscoviensis* [78]. Similar to proposed reactive oxygen species defense mechanisms in *N. defluvii* [79], all three genomes encode c type cytochrome peroxidases; Tpx, Bcp and AhpE peroxiredoxins; glutathione peroxidase gpx; thioredoxin and glutaredoxin redox proteins; methionine sulfoxide reductases MsrA and MsrB (Tables S2–S5). Unlike *N. defluvii*, all three strains encode an iron-containing superoxide dismutase (SOD) but lack catalase (CAT) gene (Table S2).

**Figure 3.**
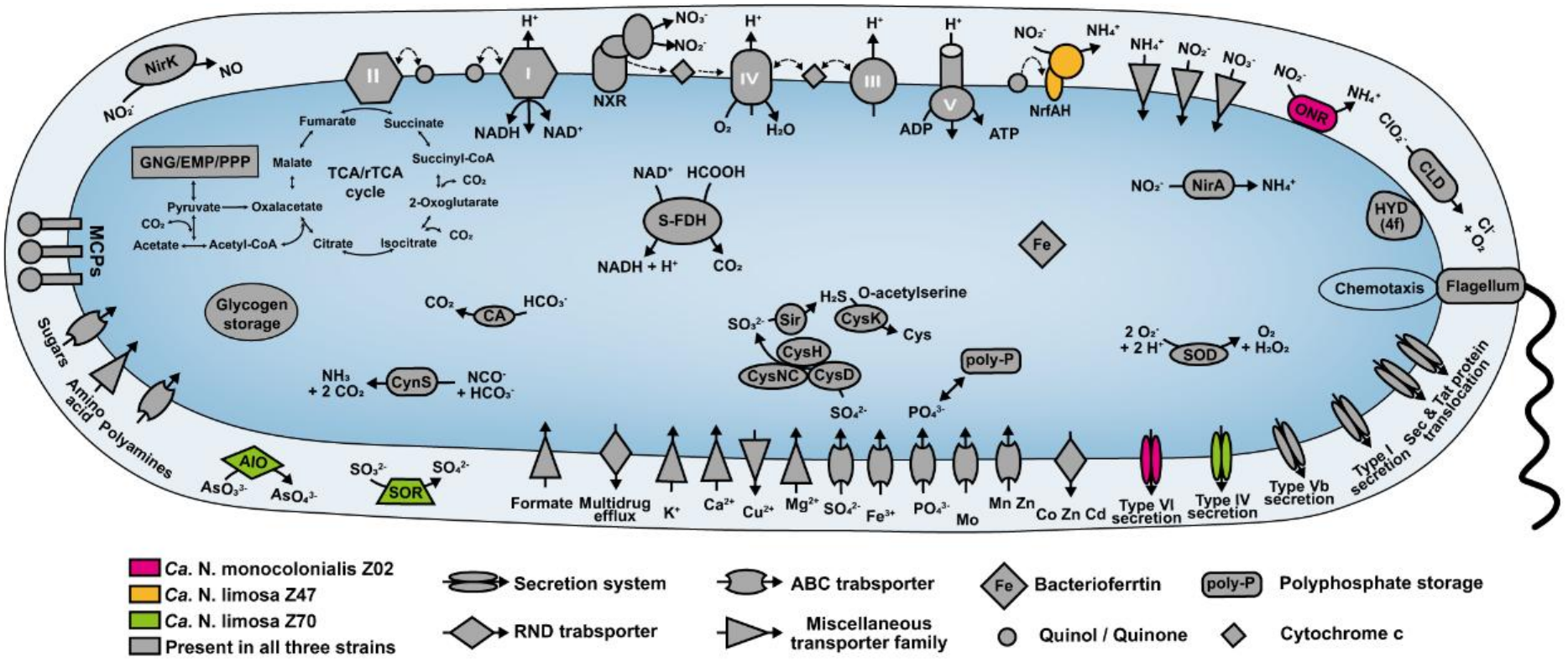
Cell metabolic cartoon constructed from the genome annotations of *Nitrospira* strains Z02, Z47 and Z70. Core functions shared by the three *Nitrospira* isolates are shown in gray, whereas features unique to strains Z02, Z47, and Z70 are shown in red, yellow, and green, respectively. NirK, nitrite reductase (NO forming); CynS, cyanate hydratase (cyanase); CA, carbonic anhydrase; S-FDH, soluble formate dehydrogenase; sir, sulfite reductase (ferredoxin); CysH, phosphoadenosine phosphosulfate reductase; CysNC, bifunctional sulfate adenylyltransferase subunit 1; CysD, sulfate adenylyltransferase subunit 2; NirA, ferredoxin-nitrite reductase; OCC, putative octaheme cytochrome c protein; SOD, superoxidase dismutase; HYD, hydrogenase; CLD, chlorite dismutase; NrfAH, cytochrome c nitrite reductase; AIO, Arsenite oxidase; SOR, cytochrome c sulfite oxidoreductase; MCPs, methyl-accepting chemotaxis proteins. Enzyme complexes of the electron transport chain are labelled by Roman numerals.

The three strains also contain some uncommon genes. For example, strain Z02 uniquely encodes a putative octaheme cytochrome c protein (OCC; AC5WRN_06445; Table S3) sharing 85.7% amino acid identity with the octaheme cytochrome c nitrite reductase (ONR) of *N. moscoviensis*, proposed to obtain electrons from the quinol pool via Pet BC for assimilatory nitrite reduction [77, 80]. A *nirA* gene (AC5WRN_06450) lies immediately downstream of the OCC encoding gene, suggesting a role in assimilatory nitrite reduction. Strain Z47 contains adjacent *nrfA* and *nrfH* genes (Table S4), as in comammox *N. inopinata*, indicating a genetic potential for nitrite reduction to ammonium. Strain Z70 contains two putative arsenite oxidase *aioBA* clusters and one putative SorAB-type sulfite: cytochrome c oxidoreductase cluster (Table S5), suggesting potential arsenite oxidation or detoxification and sulfite oxidation capacities.

### Physiological and nitrite oxidation kinetic characterizations of Nitrospira strains

Strain Z02 grew at temperature 20–37 ℃, pH 5.5–8.5, 1–21% (v/v) headspace oxygen, and 1–20 mM initial nitrite (Fig. 4A). The highest maximum specific growth rates (μ_max_) were observed at 35–37 ℃, pH 6.5, 5–10% headspace oxygen, and 1–2 mM initial nitrite. Detectable nitrite oxidation also occurred at 40 ℃ and 50 mM nitrite, but activity was too slow for complete growth curves or μ_max_ determination (Fig. S5B, C). Compared with the optimal growth rate at 5% O_2_, strain Z02 retained 78% and 85% of its rate at 1% and 21% O_2_, respectively, indicating strong oxygen adaption. Strain *N. marina* tolerates 20% oxygen and has two SOD and CAT types (Table S2) [81], whereas *N. defluvii* and related lineage I *Nitrospira* lack SOD, CAT, and superoxide reductase genes and have long been regarded as oxygen sensitive [77, 82, 83]. The three strains discovered here alter this view at both genomic and strain adaptive levels.

**Figure 4.**
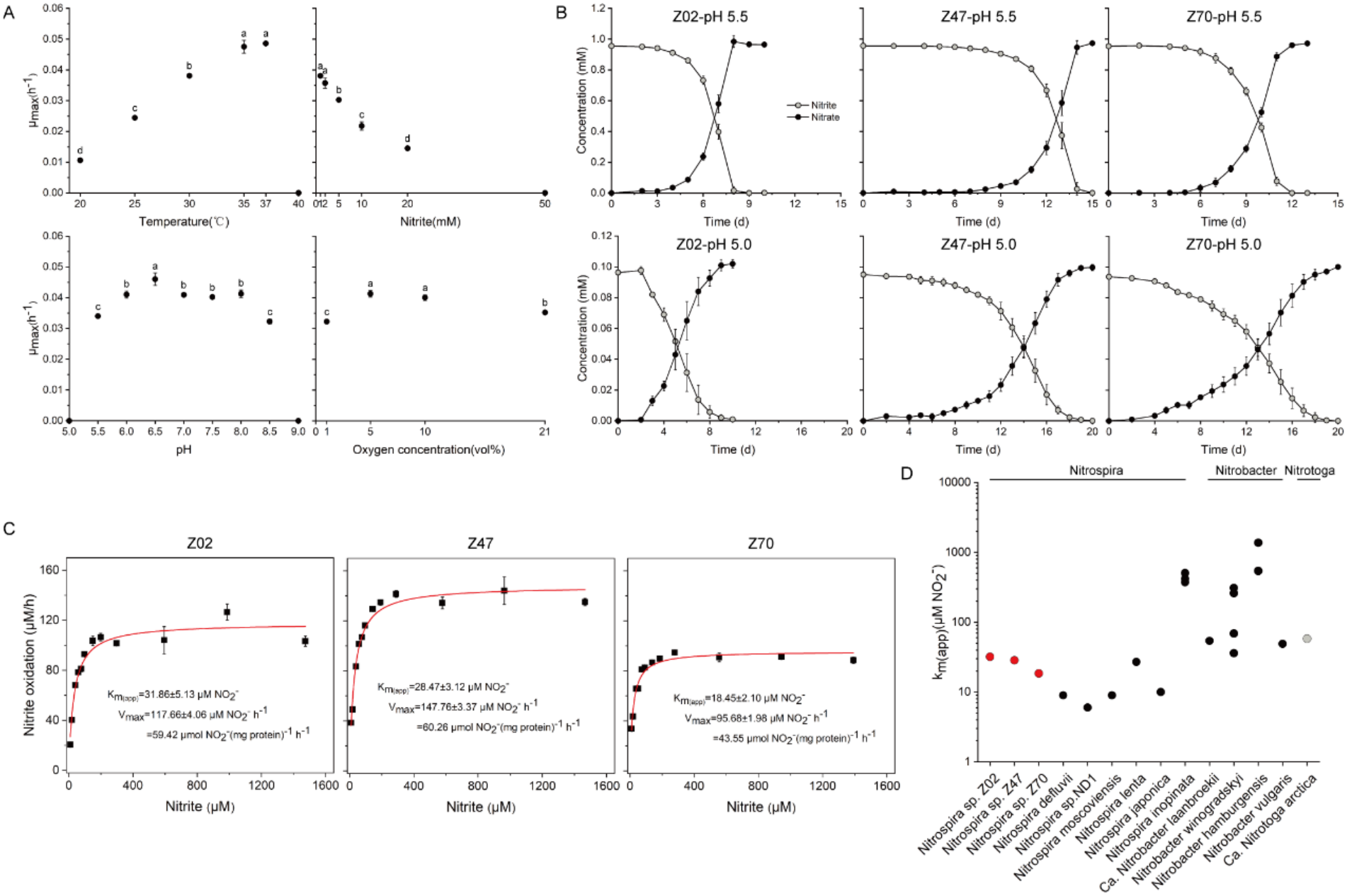
Physicochemical characterization and nitrite oxidation kinetic of *Nitrospira* strains. **A)** Apparent maximum specific growth rates (μ_max, app_) of strain Z02 estimated at different temperatures, initial nitrite concentrations, pH values, and oxygen concentrations. Different lowercase letters above the data indicate significant differences among groups (*P* < 0.05). **B)** Nitrite oxidation activity of three strains cultivated at pH 5.5 or 5.0. **C)** Michaelis–Menten kinetics of nitrite oxidation by the strains. Nitrite oxidation rates were determined from the decrease in nitrite concentration over time. The apparent half-saturation constant (K_m(app)_) and maximum nitrite oxidation rate (V_max_) were estimated by nonlinear regression using the Michaelis–Menten equation. Standard errors of the parameter estimates are reported. **D)** Comparison of K_m(app)_ values for nitrite among different NOB. The corresponding K_m(app)_ values and references are listed in Supplementary Table S3. Symbols represent the mean values of three biological replicates in panels **A** and **B** and of triplicate measurements in panels **C**. Where it is not visible, error bars are smaller than the symbols.

Compared with the optimal growth at pH 6.5, strain Z02 retained 74% of its μ_max_ at pH 5.5. All three strains grew well at pH 5.5 with 1 mM nitrite and at pH 5.0 with 0.1 mM nitrite (Fig. 4B). At pH 5.0, no activity was detected when the nitrite concentration exceeded 0.1 mM, likely due to free nitrous acid (FNA) toxicity (Table S6). These results demonstrated that the three *Nitrospira* strains are moderately acid-tolerant, plausible explaining the high abundance of *Nitrospira* in acidic environments [29–33]. Although many *Nitrobacter* strains are acid-tolerant [17–20], they are less adapted than *Nitrospira* to low oxygen and low substrate, likely making them less abundant than the *Nitrospira* in such environments [34–36].

Michaelis–Menten fitting yielded apparent nitrite half-saturation constants (K_m(app)_) of 32 ± 5, 29 ± 3 and 19 ± 2 μM for strains Z02, Z47 and Z70, with corresponding V_max_ values were 59 ± 2, 60 ± 1 and 44 ± 1 μmol NO₂⁻ mg protein⁻¹ h⁻¹, respectively (Fig. 4C; Table S7). All K_m(app)_ values were higher than that of *N. defluvii* and were closest to that of the lineage II species *N. lenta* [59] (Fig. 4D). V_max_ values of strains Z02 and Z47 were similar and exceeded the reported range for other cultivated *Nitrospira* (18–48 μmol NO₂⁻ mg protein⁻¹ h⁻¹), whereas that of strain Z70 was comparable to those of *Nitrospira* sp. ND1 and *N. defluvii* (Table S7).

### Comparative transcriptomic analysis of acid response in strain Z02

Under acidic conditions (pH 5.5) versus neutral conditions (pH 7.4), strain Z02 broadly downregulated energy metabolism and carbon fixation genes. All *nxrAB* copies and most cytochrome bd-like terminal oxidase genes were significantly downregulated: the two *nxrA* copies decreased 4.3-fold and 19.5-fold, and the cytochrome bd-like oxidase paralogue (AC5WRN_14605) decreased 1.8-fold and 2.7-fold, respectively (Fig. 5). The key carbon fixation genes, including ATP citrate lyase, fumarate reductase and several subunits of 2-oxoglutarate: ferredoxin and pyruvate: ferredoxin oxidoreductases, decreased 1.1 to 3.8-fold, and nitrogen assimilation genes decreased 2.2 to 4.8-fold decreases (Fig. S6).

**Figure 5.**
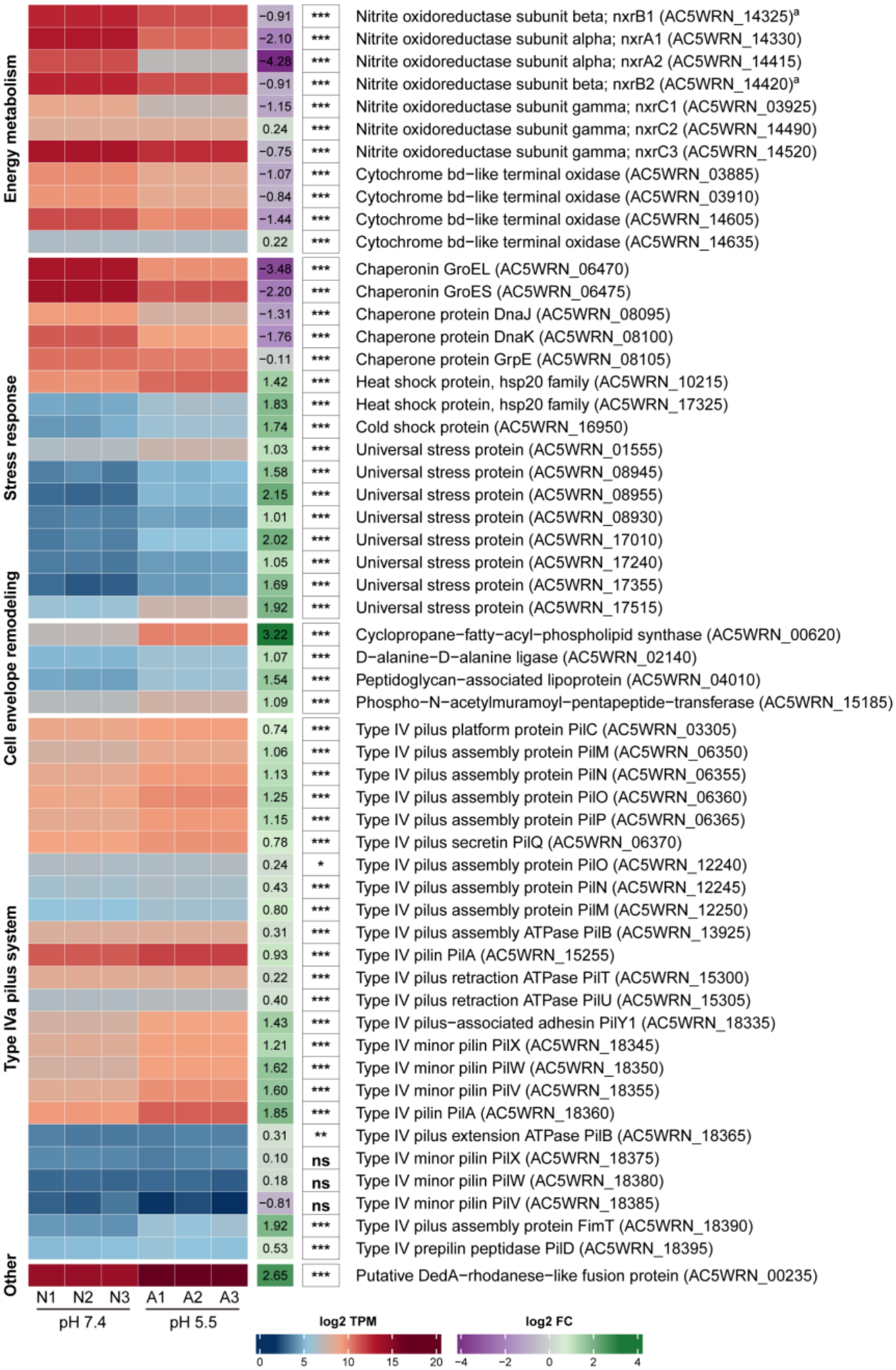
Heatmap of the differentially expressed genes in strain Z02 grown under moderately acidic and neutral conditions. Transcript abundances, expressed as transcripts per million (TPM), were log_2_-transformed. A1-A3 and N1-N3 represent three biological replicates under acidic and neutral conditions, respectively. Log_2_ fold changes (acidic relative to neutral) and adjusted P values calculated from raw counts using DESeq2 are displayed alongside each gene. Statistical significance is indicated by asterisks (adjusted P ≤ 0.001, ***; ≤ 0.01, **; ≤ 0.05, *), while not significant are denoted as “ns”. Genes are grouped by the functional categories shown on the left. Locus tags are given in parentheses. ᵃ The same estimated transcript abundance was assigned to both *nxrB* copies. Because their sequences are identical, short RNA-seq reads could not be uniquely assigned to either locus; copy-specific expression was therefore not assessed. The complete set of untransformed values are provided in Supplementary Table S8.

Acidic upregulated stress response and cell envelope remodeling genes: heat shock proteins, a cold shock protein, and multiple universal stress proteins increased 2.0 to 4.4-fold; D-alanine–D-alanine ligase, phospho-N-acetylmuramoyl-pentapeptide transferase, a peptidoglycan-associated lipoprotein and cyclopropane fatty acyl phospholipid synthase (Cfa) increased 2.1 to 9.3-fold (Fig. 5). Cfa cyclopropanates unsaturated fatty acids in membrane phospholipids; in *Escherichia coli*, membrane cyclopropanation reduces proton permeability and improves survival during acid stress [84, 85]. Notably, 23 of 24 putative type IVa pilus (T4aP)-related genes were also upregulated (Fig. 5). Two putative T4aP systems (15 and 11 genes) were identified; one contained five genes previously annotated as type II secretion systems (T2SS) and one hypothetical protein gene, all upregulated (Fig. S6, S7). Type IV pili mediate adhesion, cellular aggregation, twitching motility, microcolony development, and surface sensing [61, 86], while aggregation and biofilm growth can increase tolerance to acidic and other stresses [87, 88]. *Nitrospira* also form aggregates or flocs under changes or environmental stress [8, 25, 89]. Coordinated T4aP transcription in strain Z02 therefore probably supports a model in which stronger cell-surface interactions promote adhesion, aggregation, or biofilm-like organization under acidic conditions.

## Conclusion and outlook

To date, all known *Nitrospira* strains have been obtained mainly through dilution-to-extinction combined with related approaches (Fig. S1); none has yet been isolated by conventional single-colony isolation on common plates. Here, three lineage I *Nitrospira* strains were obtained from different environmental samples within a short period using single-colony isolation. For the first time, this study demonstrated that pure cultures of *Nitrospira* can also grow and form single colonies on solid plates made with various common gelling agents. This feasibility encourages isolation of more *Nitrospira* strains, although colony-forming ability varies among strains and unknown factors still limit growth on plates. Additionally, all three *Nitrospira* strains retained nitrite-oxidizing activity at pH 5.0, well below the pH 6.0 minimum reported for known *Nitrospira*. This is the first pure-culture demonstration that *Nitrospira* can tolerate acid conditions, like some *Nitrobacter* strains. Compared with *Nitrobacter* and other NOB, *Nitrospria* exhibit higher substrate affinity and greater low-oxygen adaptability and is therefore widely distributed and play important ecological roles in acidic environments such as acidic soils.

## Author contributions

Yueyue Zhu (Designed the research, Performed all experiments and data analyses, Prepared the manuscript), Yangyang Peng (Performed experiments and data analyses), Yuyi Zheng (Performed experiments and data analyses), Weitie Lin (Supervised the project, Reviewed the manuscript), Jianfei Luo (Supervised the project, Designed the research, Prepared the manuscript, Reviewed and edited the manuscript). All authors approved the final version.

## Data availability

The complete genome sequences of strains Z02, Z47 and Z70 have been deposited in the National Center for Biotechnology Information (NCBI) under genome accession numbers JCAJYY000000000, JCAJYZ000000000 and JCAJYX000000000 within BioProject PRJNA1480470. The corresponding raw genome sequencing reads and the RNA sequencing reads of strain Z02 are available in the Sequence Read Archive under accession numbers SRR39276770-SRR39276781.

