## Supplementary Figures for "Moderately acid-tolerant *Nitrospira* strains obtained by single-colony isolation on plates"

**Figure S1.** **Representative cultivation and isolation strategies for *Nitrospira*.**

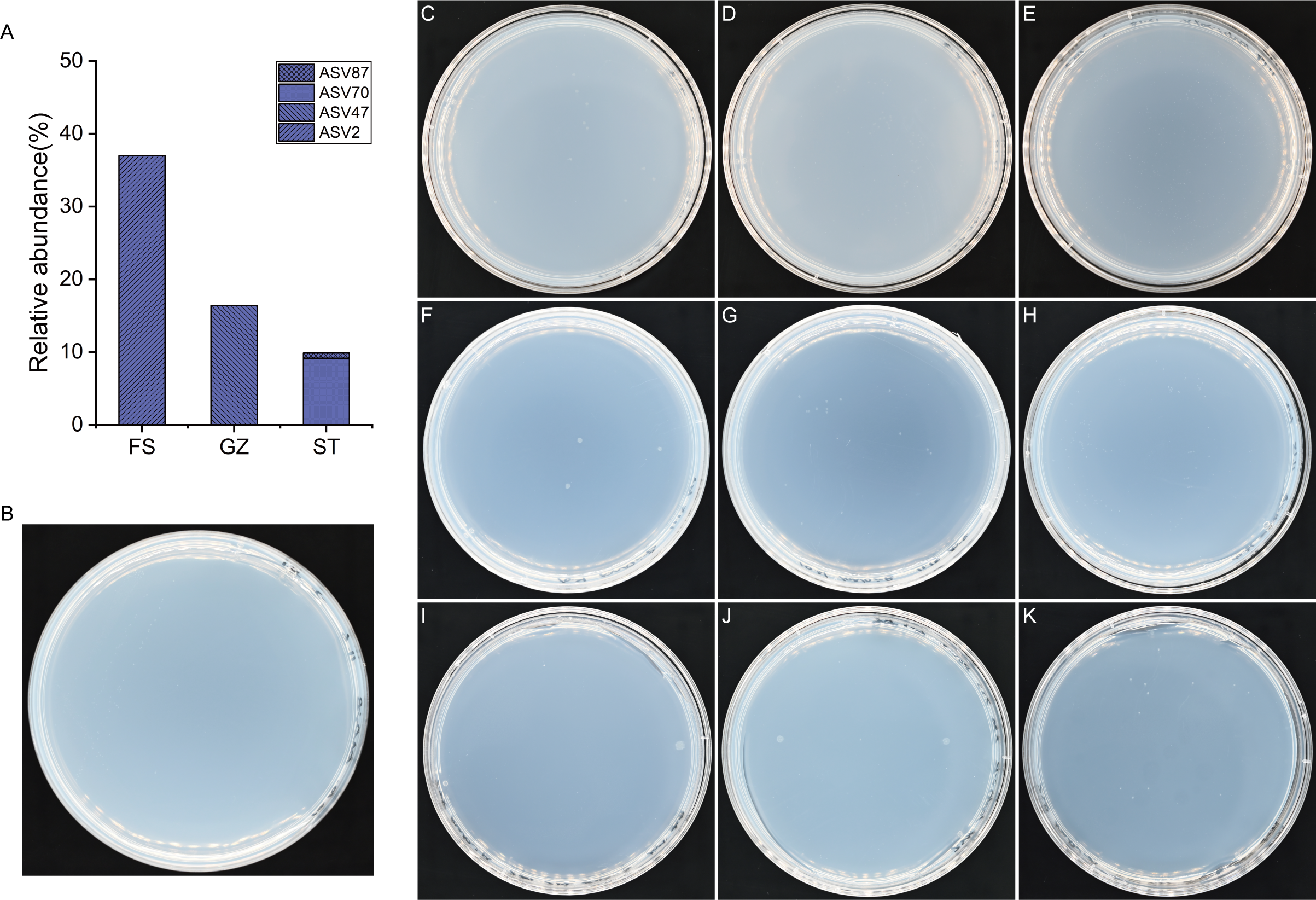

**Figure S2.** ***Nitrospira* diversity in enrichment cultures and colony growth of strain Z02. A)** ASV sequence diversity of *Nitrospira* based on full-length 16S rRNA amplicon sequencing. **B)** Colony growth of strain Z02 streak-inoculated on agarose plates after inoculation for 2 months. **C**–**K)** Colony growth of strain Z02 on agarose plates after inoculation for 8 **(C**–**E)**, 10 **(F**–**H)**, and 13 **(I**–**K)** months.

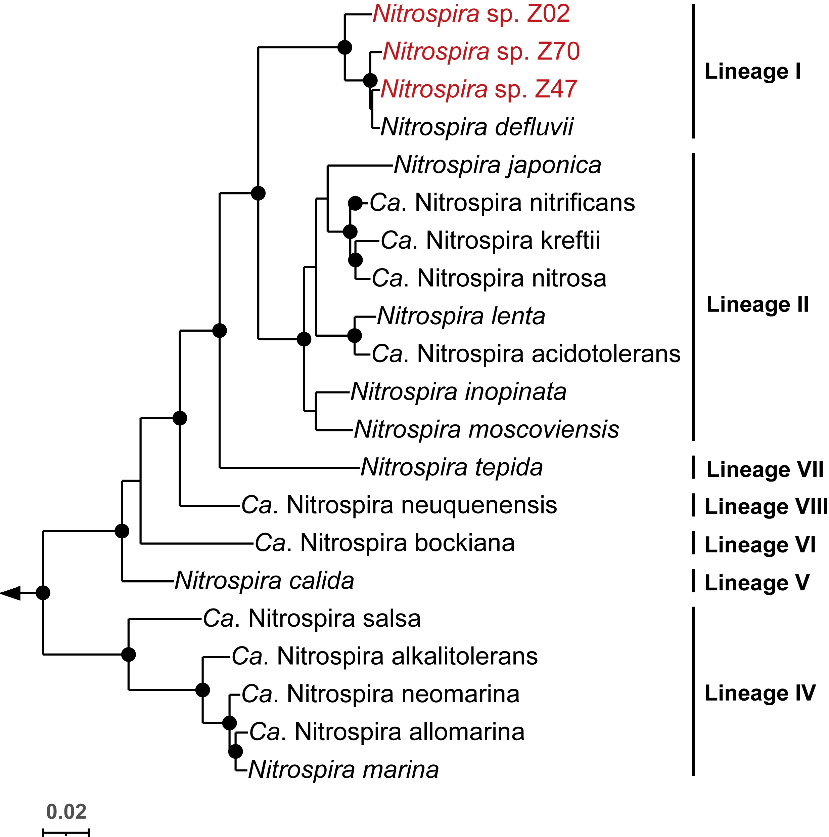

**Figure S3. Bayesian inference phylogenetic tree based on 16S rRNA gene sequences.** Full-length sequences were used, except for the 976-bp partial sequence of “*Ca*. Nitrospira neuquenensis”. A total of 1,537 nucleotide sites were retained after alignment and trimming. The tree was inferred using the GTR+Γ4 model (ASDSF = 0.0009; PSRF ≈ 1.00; minESS = 4,015). Three *Leptospirillum* 16S rRNA gene sequences from the family Nitrospiraceae were used as the outgroup, indicated by an arrow. Black dots denote branches with posterior probabilities ≥0.95. Sublineages within the genus *Nitrospira* are indicated. Strains isolated in this study are shown in red.

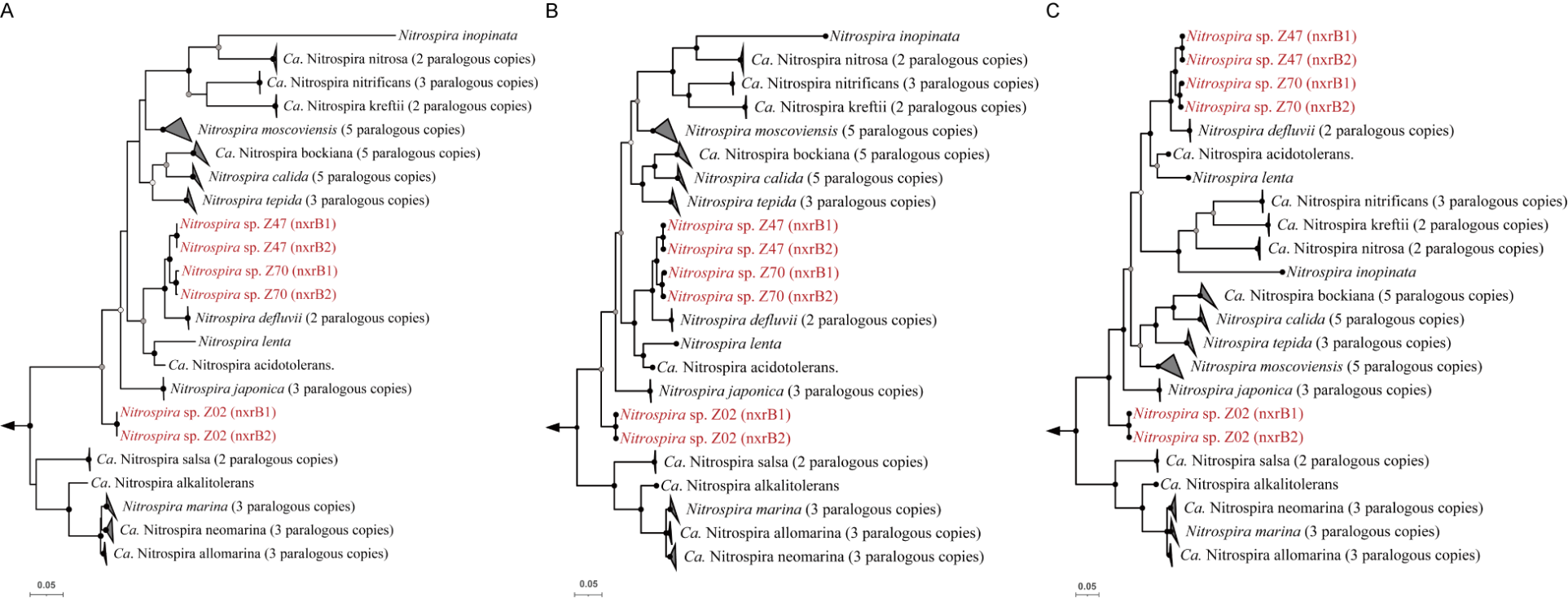

**Figure S4. Phylogenetic tree of *Nitrospira* strains based on nitrite oxidoreductase subunit B (nxrB) gene sequences.** The trees were inferred from nucleotide sequences aligned according to their amino acid translations. A total of 1,392 nucleotide sites were retained after alignment. Two paralogous nxrB copies of *Nitrospina gracilis* were used as the outgroup, indicated by an arrow. Multiple copies from the same strain are collapsed into triangles. Strains isolated in this study are shown in red. A) Maximum likelihood tree inferred using the GTR+F+I+R3 model. B) Bayesian inference tree inferred without partitioning using the GTR+I+Γ4 model (ASDSF = 0.0016; PSRF ≈ 1.00; minESS = 1,798). C) Bayesian inference tree inferred with codon position partitioning, using an independent GTR+Γ4 model for each partition and variable partition rates (ASDSF = 0.0019; PSRF ≈ 1.00; minESS = 1,272). White, grey and black dots indicate ultrafast bootstrap support values of ≥70%, ≥80% and ≥95%, respectively, in A, and posterior probabilities of ≥0.70, ≥0.80 and ≥0.95%, respectively, in B and C.

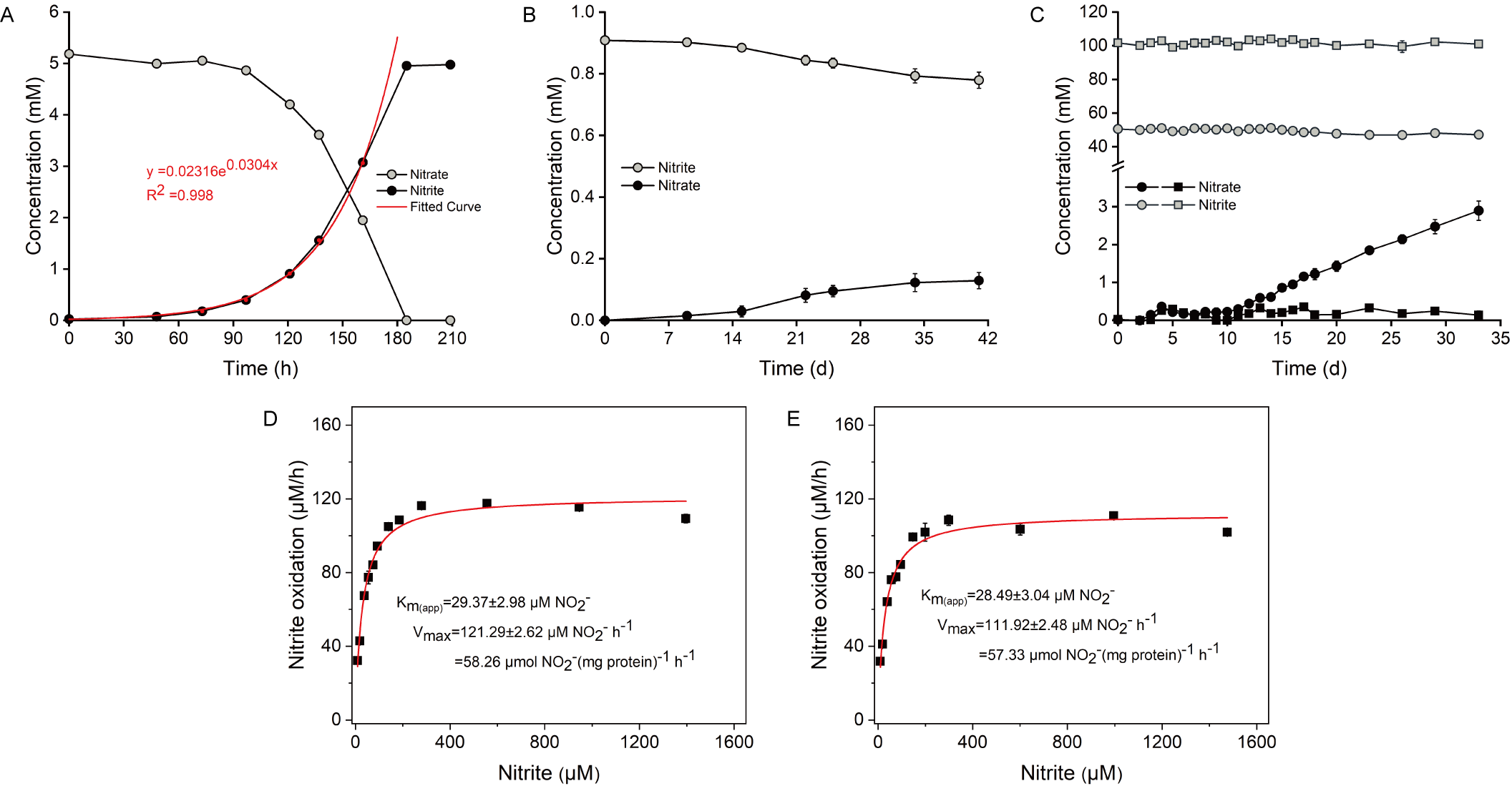

**Figure S5. Additional nitrite oxidation assays and replicate kinetic characterization of strain Z02. A)** Example of μ_max_ determination for Z02 at 5 mM initial nitrite concentration. An exponential curve was fitted to the nitrate data during the exponential growth phase (red points). The exponential parameter in the equation corresponds to the μ_max_ of the culture. **B)** Growth of Z02 at 40℃. **C)** Growth of Z02 at an initial nitrite concentration of 50 and 100 mM. **D), E)** Independent replicate measurements of Michaelis–Menten kinetics for nitrite oxidation by strain Z02. Nitrite oxidation rates were determined from the decrease in nitrite concentration over time. The apparent half-saturation constant (K_m(app)_) and maximum nitrite oxidation rate (V_max_) were estimated by nonlinear regression using the Michaelis–Menten equation. Standard errors of the parameter estimates are reported. The third biological replicate is shown in Fig. 3C. Symbols represent the means of three biological replicates in panels **B** and **C** and of triplicate measurements in panels **D** and **E**. where not visible, error bars are smaller than the symbols.

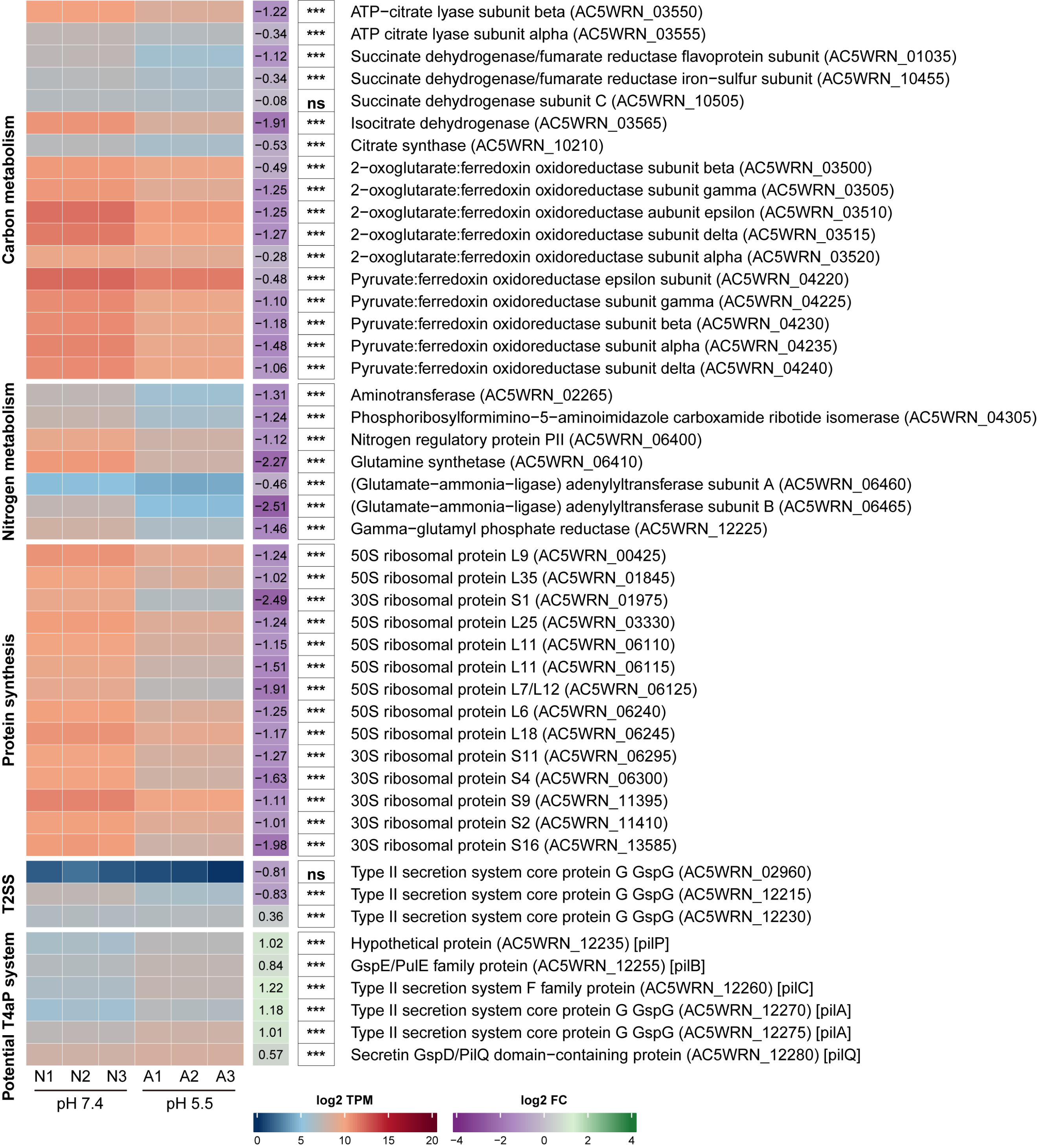

**Figure S6. Extended heatmap of additional gene expression profiles** **in strain Z02 under acidic and neutral conditions.** This figure complements Figure 5 and displays additional genes not included in the main figure. Transcript abundances, expressed as transcripts per million (TPM), were log2-transformed for heatmap visualization. A1 to A3 and N1 to N3 represent three biological replicates under acidic and neutral conditions, respectively. Log2 fold changes (acidic relative to neutral) and adjusted P values were estimated from raw count data using DESeq2 and are displayed alongside each gene. Statistical significance is indicated by asterisks (adjusted P ≤ 0.001, ***; ≤ 0.01, **; ≤ 0.05, *), while not significant are denoted as “ns”. Genes are grouped by the functional categories shown on the left. Locus tags are given in parentheses, and square brackets indicate subunits associated with Type IVa pili (T4aP) predicted by MacSyFinder. Complete untransformed TPM values are provided in Supplementary Table S8.

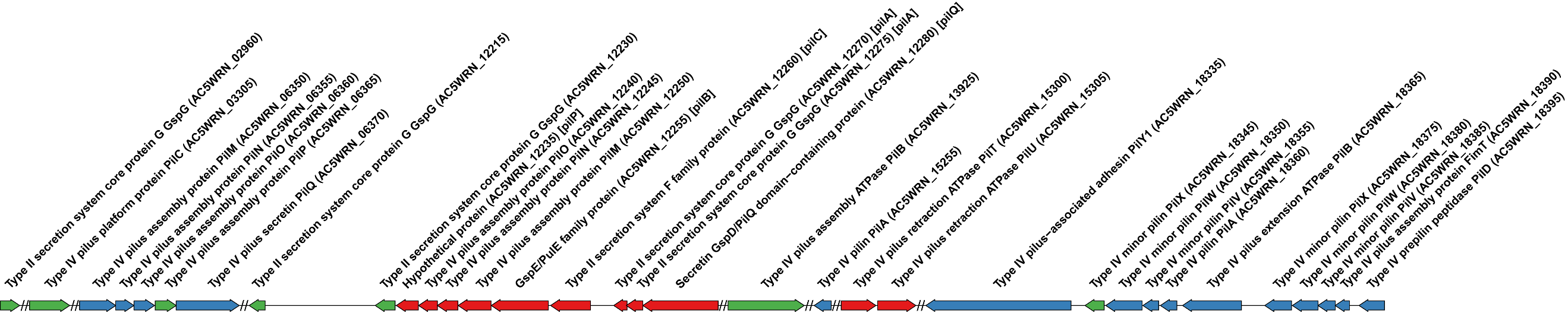

**Figure S7. Schematic representation of the genomic regions in strain Z02 that contain genes related to the Type II secretion or Type IV pilus system.** Double slashes indicate omitted genomic regions. Red and blue symbols represent genes encoding proteins associated with two independent sets of Type IVa pilus predicted by MacSyFinder. Green symbols denote genes outside the predictions. Locus tags are given in parentheses, and square brackets indicate subunits associated with Type IVa pili (T4aP) predicted by MacSyFinder.
